# Distinct temporal filters in mitral cells and external tufted cells of the olfactory bulb

**DOI:** 10.1101/135541

**Authors:** Christopher E Vaaga, Gary L Westbrook

**Affiliations:** Vollum Institute, Oregon Health and Science University, Portland OR, USA; Neuroscience Graduate Program, Oregon Health and Science University, Portland OR, USA

## Abstract

Short-term synaptic plasticity is a critical regulator of neural circuits, and largely determines how information is temporally processed. In the olfactory bulb, afferent olfactory receptor neurons respond to increasing concentrations of odorants with barrages of action potentials, and their terminals have an extraordinarily high release probability (Sicard, 1986; Murphy *et al.,* 2004). These features suggest that during naturalistic stimuli, afferent input to the olfactory bulb is subject to strong synaptic depression, presumably truncating the postsynaptic response to afferent stimuli. To examine this issue, we used single glomerular stimulation in mouse olfactory bulb slices to measure the synaptic dynamics of afferent-evoked input at physiological stimulus frequencies. In cell-attached recordings, mitral cells responded to high frequency stimulation with sustained responses, whereas external tufted cells responded transiently. Consistent with previous reports (Murphy *et al.*, 2004), olfactory nerve terminals onto both cell types had a high release probability (0.7), from a single pool of slowly recycling vesicles, indicating that the distinct responses of mitral and external tufted cells to high frequency stimulation did not originate presyaptically. Rather, distinct temporal response profiles in mitral cells and external tufted cells could be attributed to slow dendrodendritic responses in mitral cells, as blocking this slow current in mitral cells converted mitral cell responses to a transient response profile, typical of external tufted cells. Our results suggest that despite strong axodendritic synaptic depression, the balance of axodendritic and dendrodendritic circuitry in external tufted cells and mitral cells, respectively, tunes the postsynaptic responses to high frequency, naturalistic stimulation.

**Key Points:**

- The release probability of the ORN is reportedly one of the highest in the brain (Murphy et al., 2004), which is predicted to impose a transient temporal filter on postsynaptic cells.
- Mitral cells responded to high frequency ORN stimulation with sustained transmission, whereas external tufted cells responded transiently.
- The release probability of ORNs (0.7) was equivalent across mitral and external tufted cells and could be explained by a single pool of slowly recycling vesicles.
- The sustained response in mitral cells resulted from dendrodendritic amplification in mitral cells, which was blocked by NMDA and mGluR1 receptor antagonists, converting mitral cell responses to transient response profiles.
- Our results suggest that although the afferent ORN synapse shows strong synaptic depression, dendrodendritic circuitry in mitral cells produces robust amplification of brief afferent input, thus the relative strength of axodendritic and dendrodendritic input determines the postsynaptic response profile.

## Introduction

The computational capacity of neural circuits is largely determined by the short-term synaptic dynamics within the circuit (Abbott & Regehr, 2004), as determined by pre- and postsynaptic mechanisms. Short-term synaptic depression, which generally occurs at high release probability synapses, results in a net decrease in postsynaptic responses with repeated stimulation, and is often attributed to depletion of the readily releasable pool of synaptic vesicles (Liley & North, 1953; Betz, 1970; von Gersdorff & Borst, 2002; Regehr, 2012). However, at some synapses, multiple pools of synaptic vesicles with distinct release probabilities can protect the circuit from synaptic depression during high frequency stimulation (Lu & Trussell, 2016; Taschenberger *et al.*, 2016; Turecek *et al.*, 2016).

In the olfactory bulb, principal neurons receive monosynaptic input from olfactory receptor neuron afferents (Gire & Schoppa, 2009; Najac *et al.*, 2011; Gire *et al.*, 2012; Vaaga & Westbrook, 2016). Odorant receptor neurons (ORNs) respond to increasing odorant concentrations with monotonic increases in firing frequency up to 100 Hz (Sicard, 1986; Duchamp-Viret *et al.*, 1999; Rospars *et al.*, 2003; Tan *et al.*, 2010). Furthermore, the release probability of the afferent synapse between the ORN and its postsynaptic targets is one of the highest reported in the brain (ca. 0.8-0.9; Murphy *et al.*, 2004). Together, these features suggest that the transmission between ORNs and principal neurons is subject to robust short-term depression. However, *in vivo,* mitral cells respond to olfactory input with sustained responses (Giraudet *et al.*, 2002; Nagayama *et al.*, 2004; Leng *et al.*, 2014), suggesting either that release probability during trains is not as high as has been reported, or other circuit mechanisms maintain sustained transmission.

To examine the synaptic dynamics between ORN afferents and principal neurons in response to physiologically relevant stimulation frequencies, we recorded the postsynaptic responses of mitral cells and external tufted cells during high frequency afferent stimulation. Our results suggest that the high release probability and slow vesicle dynamics within the ORN are optimized for faithful transmission, but dendrodendritic amplification in mitral cells compensates for the strong synaptic depression and strongly amplifies afferent input.

## Materials and Methods

### Animals

We used adult (>p24) male and female C57Bl6/J as well as Tg(Thy1-YFP) GJrs heterozygous mice. The Oregon Health and Science University Institutional Animal Care and Use Committee approved all animal procedures.

### Slice Preparation

Olfactory bulb slices were prepared as described previously (Schoppa & Westbrook, 2001). Mice were given an intraperitoneal injection of 2% 2,2,2-tribromoethanol (0.7-0.8 mL) and monitored until fully anesthetized, then transcardially perfused with oxygenated 4 ° C modified ACSF solution, which contained (in mM): 83 NaCl, 2.5 KCl, 1 NaH_2_PO_4_, 26.2 NaHCO_3_, 22 dextrose, 72 sucrose, 0.5 CaCl_2_, 3.3 MgSO_4_ (300-310 mOsm, pH: 7.3). The brain was quickly removed and coronally blocked at the level of the striatum. Horizontal sections (300 μm) through the olfactory bulb were made using a Leica 1200S vibratome. Slices were recovered in warm (32-36° C) ACSF for 30 minutes then were stored at room temperature until transfer to the recording chamber. Unless otherwise noted, the ACSF contained (in mM): 125 NaCl, 25 NaHCO_3_, 1.25 NaH_2_PO_4_, 3 KCl, 2.5 dextrose, 2 CaCl_2_, 1 MgCl_2_ (300-310 mOsm, pH: 7.3).

### Electrophysiology

Whole cell voltage clamp and current clamp recordings were made from mitral cells and external tufted cells under DIC optics. Mitral cells and external tufted cells were identified as described previously (Hayar *et al.*, 2005; Vaaga & Westbrook, 2016). Briefly, mitral cells were identified by their soma position within the mitral cell layer and external tufted cells were identified by their relatively large soma position within the outer 2/3 of the glomerular layer. Patch pipettes (3-5 MΩ) contained (in mM): 120 K-gluconate, 20 KCl, 10 HEPES, 0.1 EGTA, 4 Mg-ATP, 0.3 Na-GTP, 0.05 Alexa-594 hydrazide, and 5 QX-314. We made no correction for the liquid junction potential (-7 mV). During cell-attached recordings, the membrane patch was held at -70 mV after achieving a gigaohm seal. Data were acquired using a Multiclamp 700b amplifier (Molecular Devices, Sunnyvale CA, USA) and AxographX acquisition software. Data was digitized at 10 kHz and low pass Bessel filtered at 4 KHz. For cell-attached recordings, the data was filtered post-hoc at 1 kHz. During whole-cell recordings the series resistance was continually monitored with a -10 mV hyperpolarizing step. Series resistance was generally <25 MΩ and was not compensated. Cells with greater than 30% change in series resistance during the recording were excluded from analysis. For better visualization, all recordings were made at 34-36 ° C.

EPSCs were elicited using single glomerulus theta stimulation, as described previously (Vaaga & Westbrook, 2016). Stimulation was provided by a constant current stimulator (100 μs, 3.2 - 32 mA) in conjunction with a small bore theta electrode (2 μm) placed directly in the axon bundle entering the target glomerulus. All recordings were made along the medial aspect of the olfactory bulb, and recordings were only made if the ORN bundle entering the target glomerulus was clearly identifiable under DIC optics. Stimulation trains (10, 25 and 50 Hz, 20 pulses) were chosen to represent the approximate firing rate of ORNs in response to odorant presentation (Sicard, 1986; Duchamp-Viret *et al.*, 1999; Carey *et al.*, 2009; Tan *et al.*, 2010). ORN stimulation was repeated at 60-second intervals, to prevent rundown. All drugs were prepared from stock solutions according to manufacturer specifications and applied via a gravity fed perfusion system. The drugs used included: 2 mM kynurenic acid, 500 nM sulpiride, 200 nM CGP55845, 10 μM CPP and 20 μM CPCCOEt. All drugs were purchased from Abcam Biochemical (Cambridge, MA, USA) or Tocris Biosciences (Ellisville, MO, USA).

### Data Analysis

Electrophysiology data was analyzed using AxographX (www.axograph.com) and IGOR Pro (version 6.22A, Wavemetrics). Spike waveforms in cell-attached recordings were detected using a threshold detection criteria in AxographX, which was used to calculate the total spike number and to generate raster plots. Voltage clamp traces represent the average of 5-10 sweeps after baseline subtraction. Fast EPSC amplitude measurements were made foot-to-peak, to eliminate any contribution of the slow current. To directly measure the slow current we recorded the EPSC amplitude just prior to each stimulus within the train. The total charge transfer (0 – 2.5 seconds after stimulus onset) was measured using a built-in AxographX routine. Data was normalized to the first fast peak EPSC amplitude, unless otherwise noted.

To estimate release probability, we used two methods to calculate the size of the readily releasable pool, each of which utilizes different assumptions (Neher, 2015; Thanawala & Regehr, 2016). In the Schneggenburger, Meyer and Neher method (SMN method), the cumulative fast EPSC amplitude (at 50 Hz stimulation) was plotted as a function of stimulus number and a linear fit was made using the last 5 responses in the train. The readily releasable pool size was estimated as the y-intercept of the linear fit (Schneggenburger *et al.*, 1999, 2002), and release probability was calculated by dividing the initial EPSC amplitude by the size of the readily releasable pool. In the Elmqvist-Quastal method (EQ method; (Elmqvist & Quastel, 1965), the fast EPSC amplitude was plotted as a function of the cumulative EPSC amplitude. A linear fit to the first 3 EPSCs was used to calculate the size of the readily releasable pool (x-intercept). Release probability was then calculated as in the SMN method.

### Statistics

All data is reported as mean±SEM unless otherwise indicated. Statistical analysis was performed in Prism6 (GraphPad Software, La Jolla, CA). One-way and two-way repeated measure experiments were analyzed using ANOVA with Holm-Sidak post-hoc pairwise comparisons as indicated in the text. To compare the exponential fit across data sets, an extra sum of squares F-test was performed to compare lines of best fit. In agreement with previous electrophysiological studies, the data was assumed to be normally distributed, and was thus analyzed using parametric statistics. Student’s paired and unpaired t-tests were used as appropriate. Sample sizes were chosen to detect an effect size of 20%, based on prior, similar experiments, with a power of 0.8. In all experiments, the initial value for α was set to p<0.05, and was adjusted for multiple comparisons as appropriate.

## Results

### Different temporal response profiles in mitral and external tufted cells

To examine the synaptic dynamics of principal neuron activity in response to high frequency afferent stimulation, we first measured the spiking of mitral and external tufted cells using cell-attached recordings. Both cell types responded to 50 Hz ORN stimulation with spikes throughout the stimulus train (Figure 1 A-D). Mitral cells and external tufted cells produced similar numbers of spikes early in the train, however, action potentials in external tufted cells gradually decreased, such that by the 7th stimulus, mitral cells produced more action potentials per successive stimulus than external tufted cells (two-way ANOVA; p<0.01; n=7 mitral cells, 8 external tufted cells; Figure 1 E). Likewise, mitral cells continued to spike well after the end of the stimulus train, contributing to the higher total number of spikes produced (mitral cells: 161.8±27.2 spikes per trial, n=7 cells; external tufted cells: 45.2±9.0 spikes per trial, n=8 cells, 2.5 second window, unpaired t-test: p=0.009, Figure 1 F).

**Figure 1:**
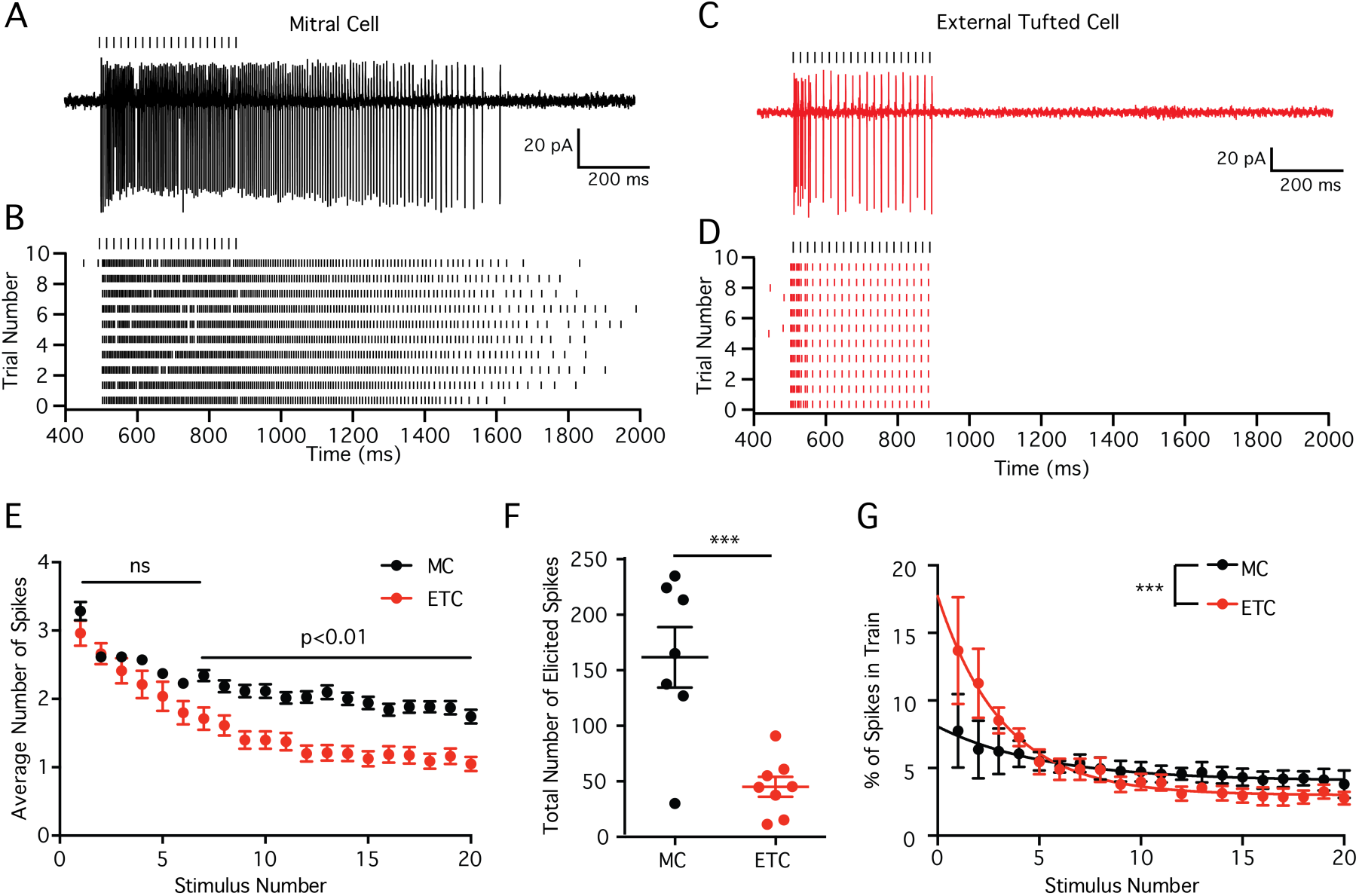
Sustained transmission in mitral and external tufted cells. (**A**) Cell attached recording from mitral cell in response to 50 Hz ORN stimulation. (**B**) Raster plot of mitral cell response. Mitral cells responded to ORN stimulation with sustained responses, which outlasted the stimulus. (**C**) Cell attached recording and (**D**) associated raster plot of external tufted cell response to 50 Hz ORN stimulation. External tufted cells produced much more transient response profiles. (**E**) Plot of the average number of action potentials produced following each stimulus in the train. Mitral cells and external tufted cells produce similar numbers of action potentials at the beginning of the train. By the end of the train, however, mitral cells produce approximately twice as many action potentials as external tufted cells. (**F**) The total number of spikes produced (within 2.5 seconds) in mitral cells is significantly higher than in external tufted cells. (**G**) Plot of the fraction of total spikes in the train as a function of stimulus number. Mitral cells (black) have a more shallow relationship, consistent with sustained transmission. External tufted cells (red) have a significantly steeper relationship, indicative of transient response profiles.

In order to quantify the temporal filter in mitral cells and external tufted cells, we calculated the percentage of the total spikes that occurred within 20 ms of each stimulus within the 50 Hz train. Using this metric, a steep input-output curve is indicative of a transient temporal filter. In both mitral cells and external tufted cells, the input-output curve was fit by a single exponential decay. In mitral cells, this relationship was relatively shallow (τ=5.2 stimuli), consistent with the observed sustained transmission. On average, mitral cells produced 7.8±2.7% of total spikes immediately after the first stimulus and 3.8±1.0% of spikes following the final stimulus (n=7 cells). In contrast, external tufted cells had a much steeper input-output relationship (τ=3.2 stimuli), producing 13.7±4.0% of total spikes after the first stimulus and 2.8±0.47% following the final stimulus (extra sum of squares F test: p<0.0001, n=8 cells). Thus the two cell types have distinct response properties with mitral cells responding to high frequency stimulation with a sustained response, whereas external tufted cells respond transiently.

### High release probability from a single pool of synaptic vesicles

Differences in release probability of ORN terminals could underlie the distinct responses of mitral cells and external tufted cells, as release probability was only examined in tufted cells (Murphy *et al*., 2004). To determine the release probability, we stimulated at high frequencies to estimate the size of the readily releasable pool using two analytical approaches as described in the methods (Elmqvist & Quastel, 1965; Schneggenburger *et al.*, 1999; Neher, 2015; Thanawala & Regehr, 2016). Consistent with a high release probability synapse, 50 Hz trains of stimuli elicited robust depression of the phasic EPSC amplitude in mitral cells (Figure 2 A_1_) and external tufted cells (Figure 2 B_1_). Both the SMN method (Figure 2 A_2_, and B_2_, C) and EQ method (Figure 2 A_3_, and B_3_, C) yielded similar estimates of the size of the readily releasable pool in mitral cells and external tufted cells. Accordingly, there was no difference in the release probability between cell types (SMN: mitral cells: 0.67±0.02, n=7 cells, external tufted cells: 0.71±0.06, n=8 cells, p=0.51; EQ: mitral cells: 0.66±0.02, external tufted cells: 0.73±0.03, p=0.14; Figure 2 C). These results indicate that the release probability of ORNs is high, but somewhat lower than previous estimates in tufted cells (Murphy *et al*., 2004), which likely reflects the activation of presynaptic D_2_ and GABA_B_ receptors in our experiments (Nickell *et al.*, 1994; Aroniadou-Anderjaska *et al.*, 2000; Ennis *et al.*, 2001; Wachowiak *et al.*, 2005; Maher & Westbrook, 2008; Shao *et al.*, 2009; Vaaga *et al.*, 2017). Consistent with this hypothesis, measurements of the release probability in D_2_ and GABA_B_ receptor antagonists (500 nM sulpiride and 200 nM CGP55845, respectively) increased the release probability to 0.95±0.06 (SMN method, n=4 cells, unpaired t-test: p=0.008). Furthermore, 2 mM kynurenic acid, which blocks receptor saturation and desensitization (Trussell *et al.*, 1993; Wadiche & Jahr, 2001; Foster *et al.*, 2002; Wong *et al.*, 2003; Chanda & Xu-Friedman, 2010) did not affect the paired pulse ratio (control: 0.24±0.05; 2 mM kynurenic acid: 0.25±0.05, n=5 cells, paired t-test: 0.70; Figure 2 D, E), suggesting that at the ORN afferent synapse, synaptic depression is primarily mediated by presynaptic factors, and is consistent with univesicular release (Murphy *et al.*, 2004; Taschenberger *et al., 2016*).

**Figure 2:**
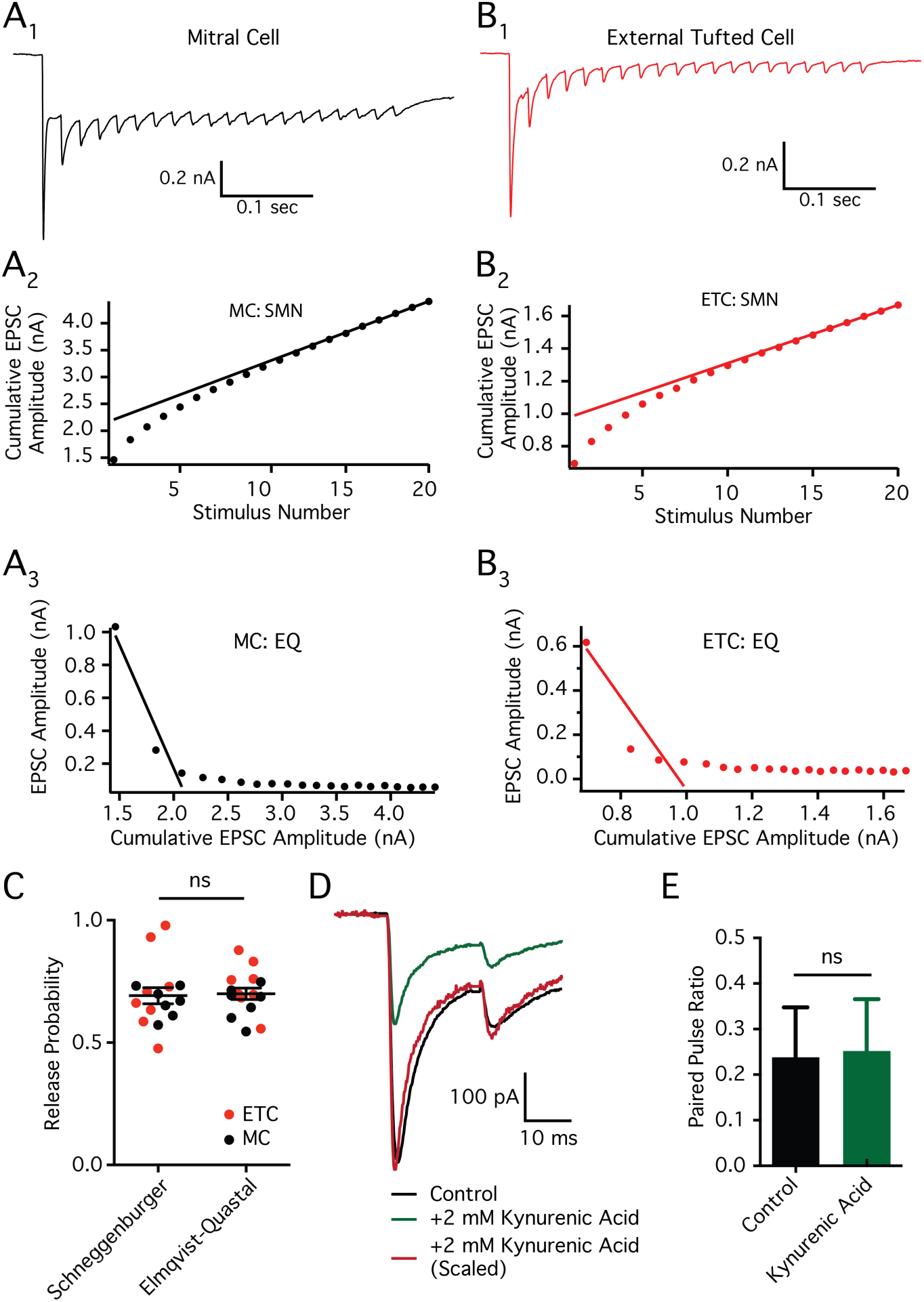
Olfactory receptor neurons have a high release probability. (**A_1_, B_1_**) Representative whole-cell voltage clamp responses to 50 Hz stimulation in mitral cells (**A_1_**, black) and external tufted cells (**B_1_**, red). (**A_2_, B_2_**) Estimates of the readily releasable pool size using the SMN train method in mitral cells (**A_2_**) and external tufted cells (**B_2_**). (**A_3_, B_3_**) Estimate of readily releasable pool size using the EQ method in mitral cells (**A_3_**) and external tufted cells (**B_3_**). (**C**) Estimates of release probability do not differ between the Schneggenburger (SMN) and Elmqvist-Quastal (EQ) methods. There was also no significant difference between the release probability calculated in mitral cells (black) and external tufted cells (red). (**D**) Paired pulse ratio in external tufted cells before (black) and after (green) addition of 2 mM kynurenic acid to prevent receptor saturation and desensitization. Response in kynurenic acid scaled to control (red). (**E**) Summary of the paired pulse ratio in external tufted cells before and after 2 mM kynurenic acid, suggesting postsynaptic saturation and desensitization do not contribute to synaptic depression.

In other circuits (Mennerick & Matthews, 1996; Sakaba & Neher, 2001; Lu & Trussell, 2016; Turecek *et al.*, 2016), multiple pools of synaptic vesicles have heterogeneous release probabilities, which, if present, could obscure our measurements of release probability and support sustained transmission at high stimulation frequencies (Neher, 2015; Turecek *et al.*, 2016). To test for the presence of multiple pools of synaptic vesicles, we stimulated at 10 Hz (20 pulses) to deplete the high release probability pool then switched to 50 Hz stimulation (20 pulses), a protocol that has been used to reveal a transient facilitation resulting from the low release probability of a separate pool of vesicles (Lu & Trussell, 2016; Turecek *et al.*, 2016). In external tufted cells this stimulation protocol failed to elicit facilitation (Figure 3 A); rather, switching to high frequency stimulation elicited further depression of the ORN-evoked phasic EPSC (EPSC_21_: 25.3±0.4% of control, EPSC_22_: 14.7±0.2% of control; Figure 3 B), suggesting a single pool of synaptic vesicles. Likewise, the decay of the phasic EPSC amplitude of external tufted cells as a function of stimulus number was best fit with a single exponential function (τ: 0.68; extra sum of squares F test: p=0.49; Figure 3 C). Together, these data indicate that a single pool of high release probability vesicles is sufficient to explain release from afferent olfactory nerve terminals.

**Figure 3:**
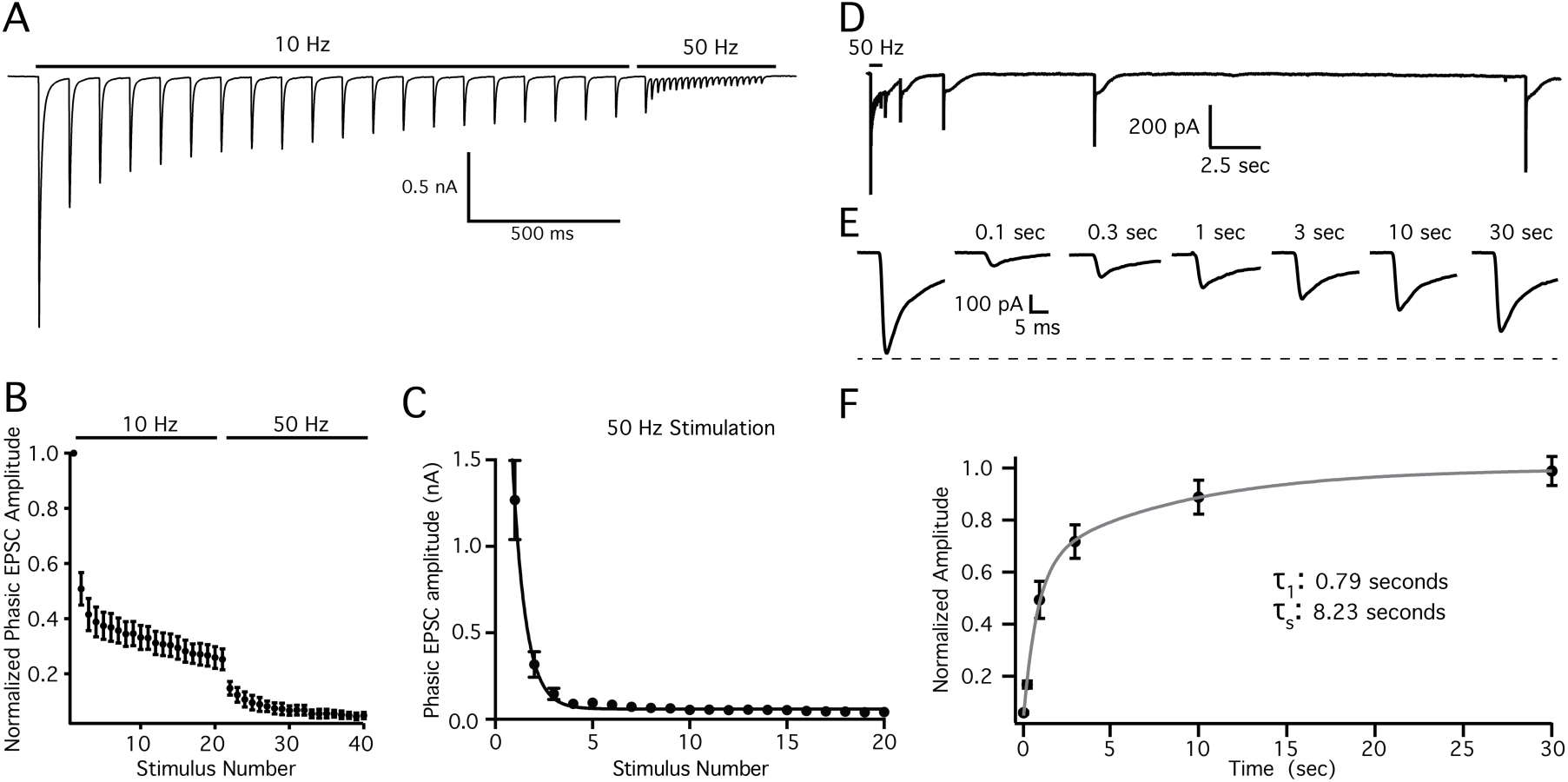
Single pool of slowly recycling vesicles. (**A**) Representative external tufted cell recording showing 10 Hz stimulation followed by 50 Hz stimulation. (**B**) Group data shows immediate depression following 10 Hz stimulation, suggesting a single pool of synaptic vesicles. (**C**) Plot of the phasic EPSC amplitude as a function of stimulus number is fit by a single exponential, further suggesting a single pool of high release probability vesicles. (**D, E**) Recovery of phasic EPSC amplitude following 50 Hz stimulation suggests that vesicle replenishment is slow. (**F**) Recovery time course is best fit by a double exponential.

Sustained responses in some cases can be maintained despite high release probability as a result of fast vesicle replenishment (Wang & Kaczmarek, 1998; Saviane & Silver, 2006). However, the phasic EPSC amplitude recovered surprisingly slowly, following a double exponential time course (τ_1_: 0.79 seconds; τ_2_: 8.23 seconds, Figure 3 D-F), suggesting that fast vesicle replenishment does not contribute to the sustained responses in mitral cells.

### Dendrodendritic excitation maintains sustained transmission

Our results suggest that properties of the afferent presynaptic terminal alone cannot explain the sustained transmission observed in mitral cells. To determine what mechanisms support sustained transmission we examined the responses of mitral cells and external tufted cells in voltage clamp following stimulation across a range of stimulus frequencies (10 Hz, 25 Hz, 50 Hz; Figure 4 A, B). Across stimulus frequencies, the phasic EPSC showed robust depression (Figure 4 D, E). Surprisingly, even relatively low stimulus frequencies (10 Hz) elicited strong depression in mitral and external tufted cells, consistent with the slow vesicle replenishment rates and unusually high release probability. In both cell types, there was a significant effect of stimulus frequency on the degree of phasic EPSC depression (One way ANOVA: mitral cell: p=0.0003; external tufted cell: p<0.0001). In both cells, the depression increased from 10 Hz to 25 Hz (mitral cells: 10 Hz: 16.4±1.3% of EPSC_1_, n=6 cells; 25 Hz: 9.4±2.1% of EPSC_1_ n=5 cells, Holm-Sidak post-hoc comparison: p<0.05; external tufted cells: 10 Hz: 14.8±2.0, n=7 cells; 25 Hz: 5.8±0.9% of control, n=7, Holm-Sidak post-hoc comparison: p<0.001), but was not significantly different between 25 Hz and 50 Hz (mitral cell: 25 Hz: 9.4±2.1% of EPSC_1_ n=5 cells, 50 Hz: 5.4±0.9% of EPSC_1_, n=6 cells, Holm-Sidak post-hoc comparison: p>0.05; external tufted cell: 25 Hz: 5.8±0.9% of EPSC_1_, n=7, 50 Hz: 4.4±0.6% of EPSC_1_, n=8 cells, Holm-Sidak post-hoc comparison: p>0.05). There was no difference in the total degree of phasic depression between mitral cells and external tufted cells at any stimulus frequency tested (Figure 4 G), consistent with similar presynaptic properties of the incoming afferents.

**Figure 4:**
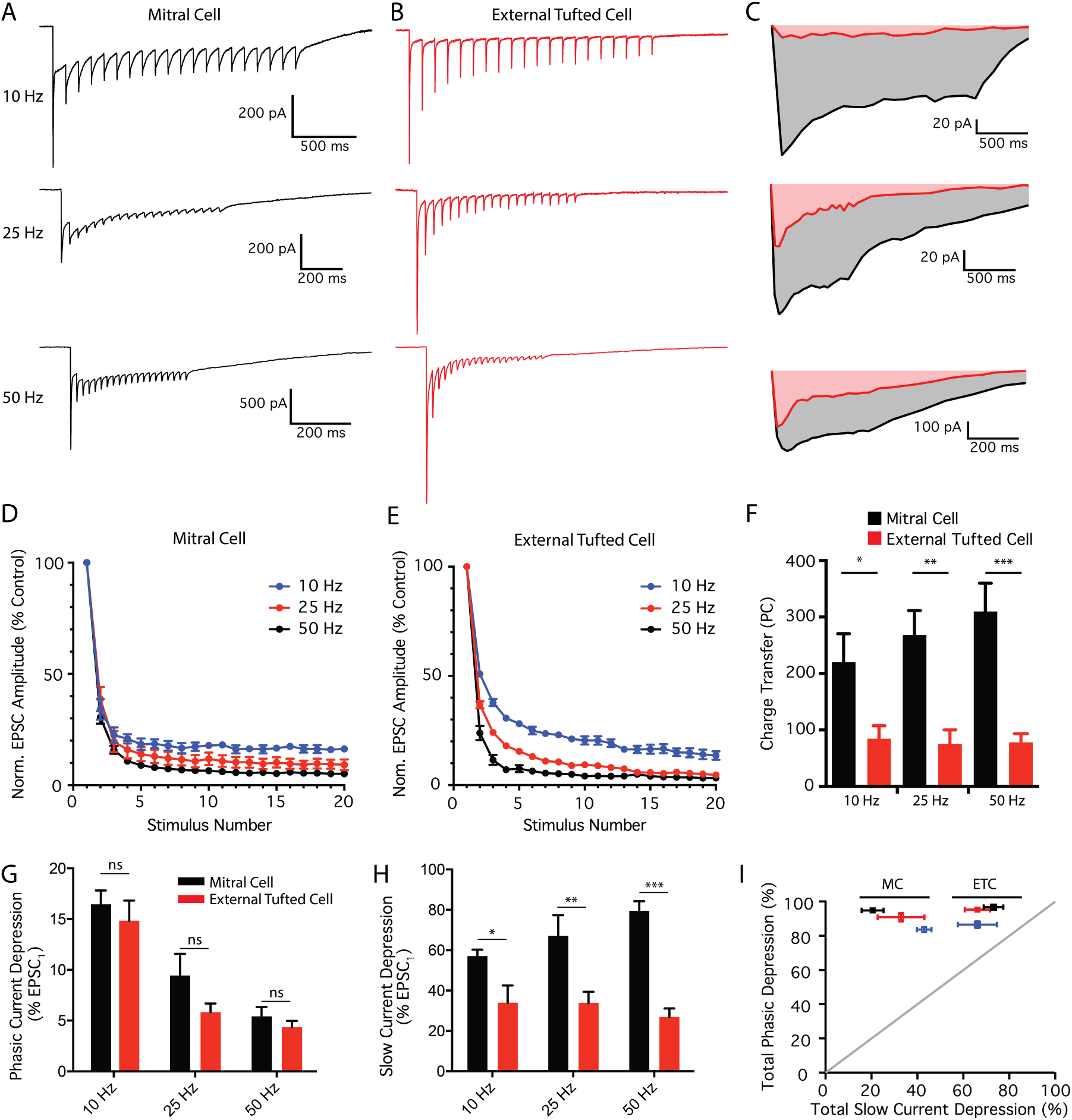
Differential modulation of phasic and slow currents in mitral and external tufted cells (A, B) Whole-cell voltage clamp responses of mitral cells (**A**, black) and external tufted cells (**B**, red) to stimulation at various frequencies (10, 25, 50 Hz). (**C**) Comparison of the slow, envelope current measured in mitral cells (grey) and external tufted cells (pink) at each stimulus frequency. Mitral cells had consistently larger envelope currents. (**D**) Depression of the phasic EPSC amplitude as a function of stimulus number in mitral cells across stimulation frequencies (blue: 10 Hz, red: 25 Hz, black: 50 Hz). (**E**) Depression of phasic EPSC amplitude as a function of stimulus number in external tufted cells (colors as in **D**). (**F**) The total charge transfer (measured 2.5 seconds after stimulus onset) was significantly larger in mitral cells than external tufted cells across all stimulation frequencies. There was no significant difference across stimulus frequencies within either cell type. (**G**) Total phasic depression in mitral cells (black) and external tufted cells (red) across stimulation frequencies. There was no significant difference between cell types at any frequency tested. (**H**) Total slow current depression in mitral cells (black) and external tufted cells (red) across stimulation frequencies. Mitral cells had significantly less slow current depression at all stimulus frequencies tested. (**I**) Plot showing a direct comparison of phasic depression and tonic depression across cell types and frequencies (blue: 10 Hz, red: 25 Hz, black: 50 Hz). Although the phasic depression was similar between cell types and frequencies, the slow current was differentially regulated in mitral cells and external tufted cells.

However in mitral cells, phasic EPSCs were superimposed on a large, slow envelope current at all stimulus frequencies, reflecting the much larger dendrodendritic currents in mitral cells compared to external tufted cells (Figure 4 C; Vaaga & Westbrook, 2016). The total charge transfer was nearly 3 times larger in mitral cells (10 Hz: mitral cell: 219.9±50.6 pC, n=6 cells, external tufted cell: 84.4±23.25 pC, n=6 cells, Holm-Sidak post-hoc comparison: p<0.05; 25 Hz: mitral cell: 268.0±43.4 pC, n=5 cells, external tufted cell: 75.5±24.7 pC, n=7 cells, Holm-Sidak post-hoc comparison: p<0.01; 50 Hz: mitral cell: 309.9±50.1 pC, n=7 cells, external tufted cell: 78.3±12.3 pC, n=7 cells; Holm-Sidak post-hoc comparison: p<0.001, Figure 4 F). Interestingly, the charge transfer was not sensitive to stimulation frequency (mitral cell: One way ANOVA: p=0.43; external tufted cell: One-way ANOVA: p=0.96, Figure 4 F), consistent with an all-or-none dendrodendritic slow EPSC (Carlson *et al.*, 2000; De Saint Jan *et al.*, 2009; Gire & Schoppa, 2009). Unlike the phasic responses, the degree of depression of the slow envelope current within the stimulus train was significantly different between mitral cells and external tufted cells (10 Hz: mitral cell: 57.1±3.2% of EPSC_1_, external tufted cell 34.0±8.4% of EPSC_1_, Holm-Sidak post-hoc comparison: p<0.05; 25 Hz: mitral cell: 67.2±10.2% of EPSC_1_, external tufted cell: 33.9±5.5% of EPSC_1_, Holm-Sidak post-hoc comparison: p<0.01; 50 Hz: mitral cell: 79.5±4.8% of EPSC_1_, external tufted cell: 26.8±4% of EPSC_1_, Holm-Sidak post-hoc comparison: p<0.0001; Figure 4 H).

In both mitral cells and external tufted cells, the depression of the phasic component was significantly larger than the depression of the slow, envelope current, and therefore all the data points fell above the unity line in a plot of phasic EPSC depression as a function of slow EPSC depression (Figure 4 I). Furthermore, the similarity of phasic depression and distinct slow current depression across cell types produced two identifiable clusters when the phasic and slow current depression are directly compared (Figure 4 I). Together this data suggests that a robust slow current supports sustained transmission in mitral cells, which is relatively insensitive to short-term depression and stimulus frequency.

### The mitral cell slow current is responsible for sustained transmission

To explicitly test the role of the slow current in generating the sustained transmission in mitral cells, we blocked NMDA and mGluR1 receptors (10 μM CPP and 20 μM CPCCOEt, respectively), which effectively blocks the slow current in mitral cells (De Saint Jan & Westbrook, 2007; Vaaga & Westbrook, 2016). As expected, bath application of NMDA and mGluR1 antagonists reduced the total charge transfer in mitral cells (mitral cell: 309.9±50.1 pC, n=7 cells; mitral cell + CPP/CPCCOEt: 42.3±8.5 pC, n=6 cells, Holm-Sidak post-hoc comparison: p<0.0001, Figure 5 A, B), to levels comparable to the charge transfer in external tufted cells (external tufted cell: 78.27±12.26, n=7 cells, Holm-Sidak post-hoc comparison: p>0.05, Figure 5 B). Thus blocking the slow current converts the mitral cell response pattern to an external tufted cell pattern.

**Figure 5:**
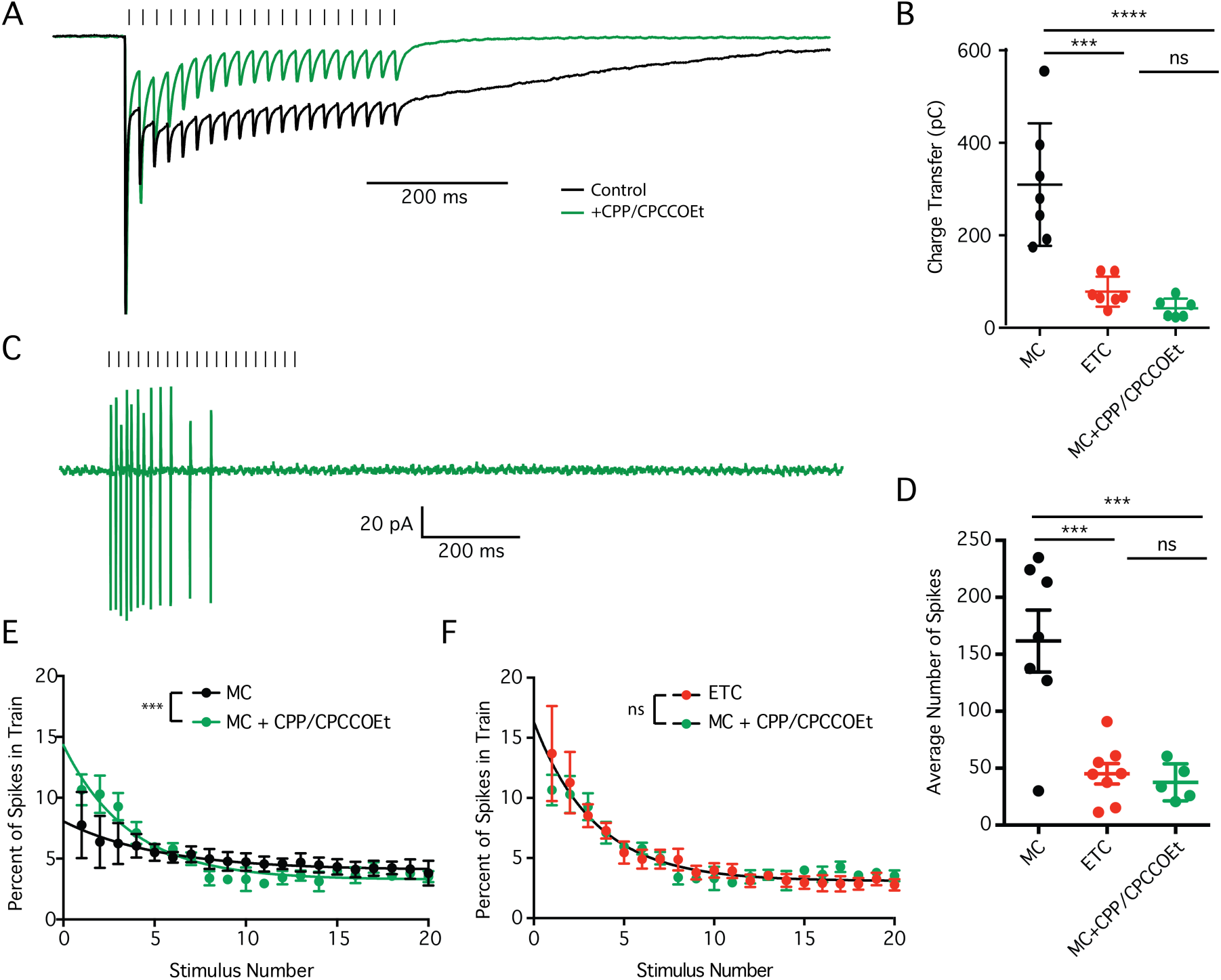
Blocking the slow current converts mitral cell responses into external tufted cell responses. (**A**) Peak scaled comparison of the whole cell voltage clamp recordings from mitral cells in control (black) and 10 μM CPP/20 μM CPCCOEt (green) in response to 50 Hz ORN stimulation. As expected, CPP/CPCCOEt blocked a significant portion of the slow envelope current. (**B**) Comparison of the total charge transfer in mitral cells (black), external tufted cells (red) and mitral cells with CPP/CPCCOEt (green) shows that blocking the NMDA/mGluR1 receptor dependent current significantly reduces the total charge transfer to levels comparable to external tufted cells. (**C**) Cell-attached recording from mitral cell in response to 50 Hz ORN stimulation shows transient spiking profile mitral cells when NMDA and mGluR1 receptors are blocked. (**D**) The total number of action potentials produced in mitral cells with NMDA and mGluR1 receptors are similar to external tufted cell responses. (**E**) Comparison of the temporal profile of mitral cell spiking in control (black) and with CPP/CPCCOEt (green). Block of NMDA and mGluR1 receptors reveal transient response profile of mitral cells. (**F**) With NMDA and mGluR1 receptors blocked, the temporal profile of mitral cell spiking (green) is not significantly different than the responses of external tufted cells (red).

In cell attached recordings of mitral cells, blocking NMDA and mGluR1 receptors also caused a 4-fold reduction in the total number of spikes produced following 50 Hz stimulation (mitral cell: 161.8±27.2 spikes, n=7 cells; mitral cell + CPP/CPCCOEt: 37.62±7.3 spikes, n=5 cells, Holm-Sidak post-hoc comparison: p<0.001; external tufted cell: 45.2±9.0 spikes, n=8 cells; Holm-Sidak post-hoc comparison: p>0.05, Figure 5 D). Furthermore, bath application of NMDA and mGluR1 receptor antagonists also altered the temporal patterning of spikes, converting the sustained responses of mitral cells to more transient responses (extra sum of squares F-test: p<0.001, Figure 5 E), which were not significantly different than the responses in external tufted cells (extra sum of squares F-test: p>0.05, Figure 5 F). Together this data suggests that differences in the amplitude of the slow current between mitral cells and external tufted cells are responsible for the sustained transmission in mitral cells, and produce their distinct temporal spiking patterns.

## Discussion

In the glomerular microcircuit, the interplay of axodendritic and dendrodendritic synapses is critical to postsynaptic processing of afferent input. Although the glomerulus has long been viewed as a cortical module whose primary function is to enhance the signal-to-noise ratio (Chen & Shepherd, 2005), the synaptic dynamics in response to high frequency, naturalistic ORN stimulation have not previously been examined. Here we demonstrate that mitral cells and external tufted cells respond to high frequency afferent input with distinct temporal filters. Mitral cells produce sustained responses, requiring dendrodendritic amplification, whereas the lack of dendrodendritic amplification in external tufted cells results in transient responses. Together, our results indicate that the axodendritic and dendrodendritic circuits are functionally separable, and the relative balance of the two circuits determines the temporal filter of the postsynaptic cell.

### Comparison with other synapses

Previous estimates of release probability using steady state measurements have suggested that the release probability of the ORN is near 1 (Murphy *et al.*, 2004). Our results using high frequency trains of stimuli suggest that the release probability of the ORN can be as high as 0.9 when presynaptic D_2_ and GABA_B_ receptors are blocked, however, in our experiments tonic and/or afferent evoked activation of presynaptic D_2_ and GABA_B_ receptors during high frequency trains reduces the release probability by approximately 30% in brain slices. Nonetheless, the presynaptic properties of olfactory receptor neurons are unusual, as compared with other synapses in the brain. Although many synapses, such as the climbing fiber synapse in the cerebellum, have a high release probability (Silver *et al.*, 1998; Dittman *et al.*, 2000), such terminals generally show multi-vesicular release (Wadiche & Jahr, 2001; Rudolph *et al.,* 2015). However, the similar paired pulse ratio in control and low affinity antagonists suggest that ORN synapses operate using univesicular release (Murphy *et al.*, 2004; Taschenberger *et al.*, 2016). Another uniquantal, high release probability synapse exists in barrel cortex between layer 4 and layer 2/3 neurons (Silver *et al.*, 2003). However, in this case, the presynaptic neuron generally fires only 1-2 action potentials in response to whisker stimulation *in vivo* (Brecht & Sakmann, 2002), so synaptic depression resulting from high release probability does not impact the postsynaptic response. The univesicular, high release probability of the ORN, therefore, is unusual because individual ORNs fires at high frequencies in response to odorants (Sicard, 1986; Duchamp-Viret *et al.*, 1999; Carey *et al.*, 2009; Tan *et al.*, 2010).

In theory, the high frequency transmission of the ORN could be maintained despite a high initial release probability through multiple mechanisms including fast vesicle recycling (Kushmerick *et al.*, 2006; Saviane & Silver, 2006) and a second pool of low release probability vesicles (Lu & Trussell, 2016; Taschenberger *et al.*, 2016; Turecek *et al.*, 2016). These properties, however, appear to be absent in ORNs. In fact, the recovery of the fast EPSC following depletion was approximately 10 fold slower than at the calyx of Held (Kushmerick *et al.*, 2006), suggesting that individual ORNs may only transiently contribute to the postsynaptic response, thereby providing a rationale for the massive convergence of unimodal ORNs onto single glomeruli.

### Axodendritic input is tuned to ensure faithful transmission

A striking feature of the glomerular microcircuit is the massive convergence of axons to a single glomerulus, with each axon carrying functionally redundant information (Mombaerts *et al.*, 1996). This unimodal input is critical for odorant identification, as each odorant mixture elicits a unique map of activated glomeruli, a so-called odor image (Xu *et al.*, 2000). However, from a computational perspective, the massive redundancy is a waste of information channels (Rieke, 1999; Chen & Shepherd, 2005). Furthermore, each olfactory receptor neuron responds to increases in odorant concentration with monotonic increases in firing frequency, reaching up to 100 Hz (Sicard, 1986; Duchamp-Viret *et al.*, 1999; Carey *et al.*, 2009; Tan *et al.*, 2010). Coupled with the high ORN release probability (Murphy *et al.*, 2004), trains of ORN activity produce strong synaptic depression as demonstrated in our experiments, imposing a transient temporal filter in postsynaptic cells, resulting from presynaptic vesicle depletion.

An advantage of such a high initial release probability is that odorant binding events in the periphery are faithfully transmitted to the olfactory bulb in a nearly all-or-none manner. The olfactory system is exquisitely sensitive, capable of detecting odorants at concentrations as low as 1 part per 10^15^ molecules (Julius & Katz, 2004). In the periphery, this high sensitivity is achieved through biochemical amplification downstream of G-protein coupled odorant receptors, such that a single odorant receptor-binding event can elicit an action potential in the ORN (Lynch & Barry, 1989). The high release probability of ORNs maintains the high sensitivity of the olfactory system, by ensuring that ORN activity is faithfully converted to a postsynaptic response. However, this circuit design comes at a cost in that individual nerve terminals can only transiently contribute to postsynaptic activation, therefore requiring an ensemble of functionally redundant channels to accurately convey information with high fidelity.

### Dendrodendritic circuitry promotes sustained transmission

The high release probability of axodendritic input comes at another cost: the “noisy” olfactory environment dramatically increases the total number of activated glomeruli in response to ambient air. The signal to noise ratio, therefore, is enhanced through multiple mechanisms, including the dendrodendritic amplification within the glomerulus (Carlson *et al.*, 2000; Chen & Shepherd, 2005; De Saint Jan & Westbrook, 2007; Vaaga & Westbrook, 2016). The robust increase in synaptic charge associated with the slow, dendrodendritic current effectively converts the transient axodendritic input into a sustained spiking response, greatly amplifying afferent input. Interestingly, our results indicate that only a subset of excitatory neurons, the mitral cells, within the glomerulus express dendrodendritic amplification (Vaaga & Westbrook, 2016).

### Parallel input paths convey temporally distinct information

Different principal neuron subtypes in the olfactory bulb represent parallel input pathways. For example, *in vivo*, tufted cells respond to lower odorant concentrations, have concentration invariant responses, and respond to odorants earlier in the sniff cycle (Nagayama *et al.*, 2004; Igarashi *et al.*, 2012; Fukunaga *et al.*, 2012; Kikuta *et al.*, 2013). Mitral cells, on the other hand, are more narrowly tuned than tufted cells, and shift their responses relative to the sniff cycle in response to increasing odorant concentrations (Nagayama *et al.*, 2004; Kikuta *et al.*, 2013). These *in vivo* results are consistent with the view that tufted cell responses maintain the sensitivity of the ORN, via strong afferent evoked responses, whereas mitral cells provide robust amplification, via strong dendrodendritic circuitry.

Recent evidence suggests that within piriform cortex the concentration invariant network of activated pyramidal cells encodes odorant identity whereas concentration is encoded by the temporal response profiles of pyramidal cells (Bolding & Franks, 2017). More specifically, the spiking patterns of these pyramidal cells have two distinct peaks, one with a short latency and one with a longer latency. As concentration increases, the relative lag between these two responses is shortened (Bolding & Franks, 2017). Mechanistically, this may result from the integration of olfactory bulb projection neurons that express strong axodendritic input, contributing to the short latency, concentration invariant response, and neurons that express strong dendrodendritic input, with variable, concentration-dependent delays. Such an activation scheme, however, would require overlapping projection patterns in piriform cortex. Single axon tracing studies, however, suggest that mitral cells and tufted cells project to largely non-overlapping regions of olfactory cortex (Igarahsi *et al*., 2012). More specifically, mitral cells project to dorsal region of the anterior piriform cortex as well as the cortical region of the olfactory tubercle, posterior piriform cortex and tenia tecta; whereas tufted cells, including external tufted cells, project to the ventrorostral anterior piriform cortex, the cap of the olfactory tubercle, and the pars extrema and the poteroventral region of the anterior olfactory nucleus. (Igarashi *et al.*, 2012). Resolving the exact projection patterns and mechanisms behind generating distinct timing signals in piriform cortex is critical to understanding the encoding of concentration within the olfactory system. Our results, however, demonstrate that the distinct balance of axodendritic and dendrodendritic synaptic strength in each principal cell population likely contributes to the unique computations within these parallel input pathways, by imposing unique temporal filters in each cell type.

## Acknowledgements

We thank Dr. Henrique von Gersdorff and members of the Westbrook lab for helpful comments on this manuscript. This work was supported by a NS26494 (GLW, a National Science Foundation Graduate Research Fellowship DGE 0925180 (CEV), and a LaCroute Neurobiology of Disease fellowship (CEV).

